# Demographic variability and heterogeneity among individuals within and among clonal bacteria strains

**DOI:** 10.1101/105353

**Authors:** Lionel Jouvet, Alexandro Rodríguez-Rojas, Ulrich K. Steiner

## Abstract

Identifying what drives individual heterogeneity has been of long interest to ecologists, evolutionary biologists and biodemographers, because only such identification provides deeper understanding of ecological and evolutionary population dynamics. In natural populations one is challenged to accurately decompose the drivers of heterogeneity among individuals as genetically fixed or selectively neutral. Rather than working on wild populations we present here data from a simple bacterial system in the lab, *Escherichia coli.* Our system, based on cutting-edge microfluidic techniques, provides high control over the genotype and the environment. It therefore allows to unambiguously decompose and quantify fixed genetic variability and dynamic stochastic variability among individuals. We show that within clonal individual variability (dynamic heterogeneity) in lifespan and lifetime reproduction is dominating at about 82-88%, over the 12-18% genetically (adaptive fixed) driven differences. The genetic differences among the clonal strains still lead to substantial variability in population growth rates (fitness), but, as well understood based on foundational work in population genetics, the within strain neutral variability slows adaptive change, by enhancing genetic drift, and lowering overall population growth. We also revealed a surprising diversity in senescence patterns among the clonal strains, which indicates diverse underlying cell-intrinsic processes that shape these demographic patterns. Such diversity is surprising since all cells belong to the same bacteria species, *E. coli*, and still exhibit patterns such as classical senescence, non-senescence, or negative senescence. We end by discussing whether similar levels of non-genetic variability might be detected in other systems and close by stating the open questions how such heterogeneity is maintained, how it has evolved, and whether it is adaptive.

**Data deposition:** The processed image analysis data, R code, as well as the Leslie matrices will be archived at Dryad.org.

---

Heterogeneity among individuals has important ecological and evolutionary implications because it determines the pace of ecological and evolutionary adaptation and shapes eco-evolutionary feedbacks (Hartl and Clark 2007, Steiner and Tuljapurkar 2012, Vindenes and Langangen 2015). Despite substantial methodological and empirical efforts, it remains challenging to unambiguously differentiate the causes that drive the observed heterogeneity among individuals in their life courses, their traits, and their fitness components (Steiner and Tuljapurkar 2012, Bonnet and Postma 2016, Cam et al. 2016). There is consensus that heterogeneity among individuals is caused by changes in the environment, by variation in the genotype, by the genotype-by-environment interaction, and by noise or intrinsic processes many of which show stochastic properties (Endler 1986, Finch and Kirkwood 2000, Kirkwood et al. 2005). The latter cause has either been deemed as noise associated with non-biological processes, e.g. measurement error, and with unknown hidden processes that were of little biological relevance. Alternatively, this intrinsic “noise” has been investigated for underlying biological processes with stochastic characteristics and its substantial biological implications are illustrated by quantitative genetic and population genetic studies. The interest in such intrinsic noise is best understood by its slowing of evolutionary dynamics via lowering heritabilities and enhancing genetic drift (Lande et al. 2003, Hartl and Clark 2007).

The challenge is heightened in natural populations to decompose the observed heterogeneity into its genetic, environmental, and non-genetic, non-environmental — stochastic — component. In such populations, we are confronted with high genetic diversity and complex environmental and gene-by-environment interactions (Fitzpatrick et al. 2016). The knowledge about the genotypes at the individual level is limited (e.g. pedigree) or in many cases totally absent. Certain environmental variables are known at the population level, but micro-environmental differences are less explored. The response of individuals to the known population level environmental factors varies — e.g. due to gene-by-environment interactions — and individuals are differently affected by the population level environment, e.g. not all individuals are exposed equally. Ecologists agree on that one cannot encompass the whole complexity of natural systems and hence the additional variance is a combination of error and some hidden drivers of heterogeneity. The aim remains identifying the cause of this additional heterogeneity since only such identification allows forecasting of and understanding of evolutionary and ecological population dynamic processes (Lande et al. 2003, Tuljapurkar et al. 2009, Steiner et al. 2010).

Not only empirical challenges occur when trying to decompose the observed variance in natural populations, from a methodological point of view challenges await us. Various statistical approaches aim at classifying the hidden heterogeneity as either fixed at birth, e.g. additive genetic effects or maternal effects, or as dynamic heterogeneity, heterogeneity generated during the course of life (Tuljapurkar et al. 2009, Steiner et al. 2010, Steiner and Tuljapurkar 2012, Bonnet and Postma 2016, Cam et al. 2016, Hartemink et al. 2017). Such models, be they based on mixed effect models, Markov chains, hidden Markov chains, covariate models or related models, are biased and cannot reveal the accurate underlying mechanism unless the contributing factors and the underlying error structure is known (Bonnet and Postma 2016, Cam et al. 2016). This applies to both so-called neutral models that base their arguments on dynamic heterogeneity — heterogeneity best described by stochastic transitions among stages that shape individual life courses —, and adaptive selective models that base their arguments on fixed heterogeneity, variability among individuals fixed at birth described by genetic differences or maternal effects. Note, in the fixed type of models there remains a large unexplained residual error, a variance of unknown origin, and even models that combine dynamic and fixed heterogeneity suffer from biased estimations.

To circumvent these empirical and methodological challenges faced in natural populations, we used here cutting-edge microfluidic technologies on a simple bacterial system in the lab, *Escherichia coli.* This system, in combination with age-structured matrix population models, allowed us to unambiguously decompose and quantify fixed, genetic variability and dynamic, stochastic variability among individuals. The highly-controlled environment of the microfluidic system excluded extrinsic environmental variation and gene-by-environment variation, sharpening the focus on decomposing genetic and non-genetic and non-environmental individual variability. We defined all genetic variability as the variance among seven (clonal) bacteria strains in their mean fitness components. We call this among strain genetic variability in fitness components fixed heterogeneity. This fixed heterogeneity is set in relation to dynamic heterogeneity, the variability in fitness components among individuals within strains. This dynamic heterogeneity is generated by cell intrinsic processes and can be best described as neutral individual heterogeneity. At least it is non-genetic and non-environmental induced variability, and demographic characteristics are not heritable between mother and daughter cells (Steiner and Tuljapurkar 2012, Steiner et al. 2017). As expected, fixed, among strain genetic variability was modest compared to the substantial within strain variability in reproduction and survival. Variance in lifespan within strains explained ~88% and 12% was related to among strain variance in lifespan. Variance in lifetime reproductive success within strains explained ~82% and 18% was related to among strain variance. Our finding does not imply that the genetic variability is not relevant, it just highlights that there is large amount of heterogeneity expressed within strains that is neither genetically nor environmentally driven and can therefore be described as neutral.

## Material and methods

The study organism we worked with is *E. coli*, a rod-shaped bacteria and molecularly well explored model organism. We defined each bacteria cell as an individual. Individuals grow (elongate) and reproduce by binary fission, a division in two usually equal sized cells. Note that these cells are functionally unequal, and such functional asymmetry is crucial otherwise a mother cell would divide into two identical daughter cells and thereby the original mother cell would “die” (Johnson and Mangel 2006, Tyedmers et al. 2010). In addition, populations with perfect symmetric dividing cells are not viable over multiple generations if oxidative damage accumulates in cells as described for many aging processes including those for bacteria (Ackermann et al. 2007, Evans and Steinsaltz 2007, Lindner and Demarez 2009, Tyedmers et al. 2010). The asymmetry in division allows to distinguish a mother cell, the cell that holds the old pole of the cell wall, and a daughter cell, the offspring cell that inherits the more recent pole of the cell wall (Stewart et al. 2005). Even though senescence patterns are observed and individual cells age, the change in mortality rates across age is not determined by the age of the cell pole itself even though it is correlated(Steiner et al. 2017). It is not the cell pole age, but the cytoplasm content that influences mortality rates. While the mother cells senesce, the daughter cells are thought to be rejuvenated (Ackermann et al. 2007), but this rejuvenation seems only to be perfect for daughters of young mothers and not for daughters of old mothers (Steiner et al. 2017). Further, among isogenic bacteria the lifespan of the mother does not correlate with the lifespan of the daughter, which suggests that the asymmetry at fission has a dominating stochastic component to it (Steiner et al. 2017). Despite intensive mechanistic research on the factors involved in the functional asymmetry, none of the factors have been identified as the actual cause or consequence of the functional difference that determine the cell fates (Nyström et al. 2007, Lindner and Demarez 2009, Tyedmers et al. 2010). The more quantitative demographic approach we have taken here does not focus on the within cell mechanistic factors but rather aims at decomposing the genetic and dynamics components driving individual heterogeneity.

For our experiments we used a bacterial microfluidic system called mother machine (Wang et al. 2010, Steiner et al. 2017) (Fig.1). This system allows tracking thousands of individual cells via time-lapse phase-contrast microscopic imaging. Using these time-lapse images, we determined for each (mother) cell the lifespan, the timing and number of divisions, as well as the size and cell elongation throughout their lives. We identified cell death by propidium iodide, a chemical that enters the cell after the cell wall lysed and emits a strong red fluorescent signal when it binds to the DNA. We only collected and tracked data on the (old pole) mother cell, the bottom most cell of the dead-end side channels (Fig. 1); daughter cells are pushed out into the main, laminar flow channel and washed away and cannot be tracked throughout their lives.

**Fig. 1:**
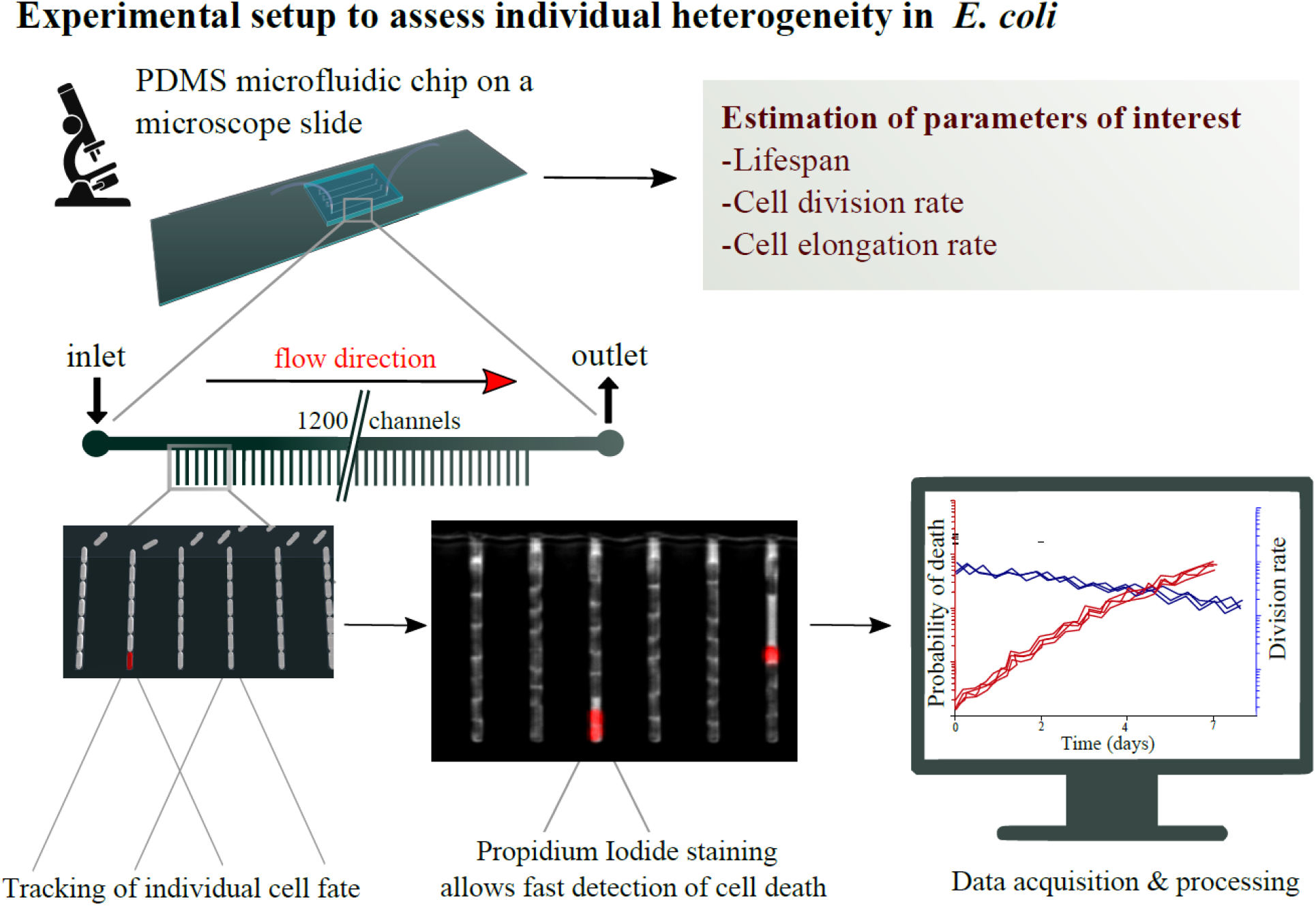
Experimental setup of microfluidic system to track individual bacteria cells throughout their lifetime. The main (horizontal) channel with the constant laminar flow connects directly the inlet and outlet and provides the cells, that grow in the vertical side channels, with fresh media. The vertical smaller side channels hold at their dead end the focal (mother) cell, which is the bottom most cell in each side channel.

To control the genetic variability, we conducted separate experiments for seven different isogenic strains. Based on the individual demographic data of the tracked cells, we estimated hourly age-specific survival and reproduction rates which we used to parameterize age-specific matrix models (Leslie matrices), one model for each of the seven isogenic strains. We selected the seven *E. coli* strains based on their common use as model organisms in the lab (K12 variants) and complemented them by a genetically distinct strain (Fig. 2). Details on the experiments, microfluidic chip production and strains are given in the online Supplementary material.

**Fig. 2:**
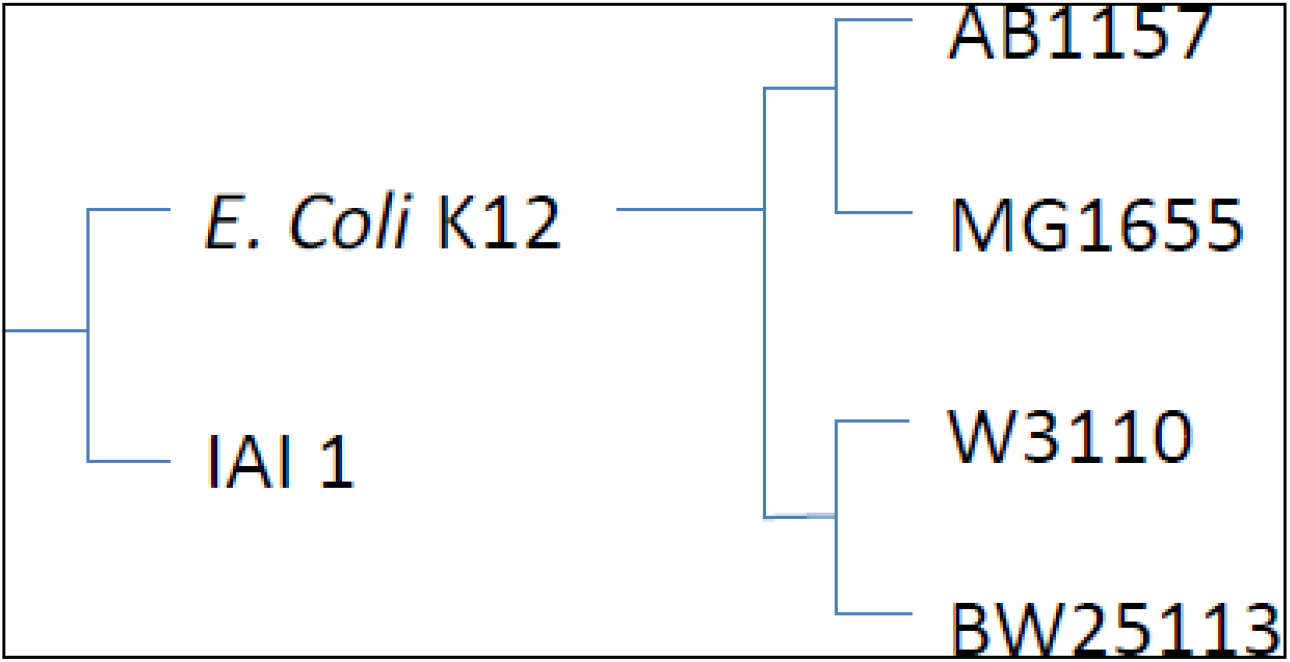
Phylogenetic relationship among the different *E. coli* strains.

### Data analysis

To analyse our individual level demographic data (each mother cell is an individual) that was collected by time-lapse imaging, we parameterised discrete age-structured population models formulated as Leslie matrices. The time-lapse images were taken at 4-minute intervals, and therefore we recorded for each focal (mother) cell whether it had divided, how much it had grown, and whether it died within a 4-minute time interval. For each isogenic strain, we formulated one Leslie matrix, **A** (Table 1). Since not all cells were dead at the end of the experiments we right censored these cells. To parameterise the age-structured Leslie matrix models we calculated hourly division and survival rates rather than 4-minute rates as collected via the time-lapse imaging. Hourly rates were calculated to reduce uncertainly due to sampling variability of small sample sizes and to increase the accuracy of calculating vital rates. To be precise, for each strain, we calculated the age-specific survival probabilities from time *t* to time *t+1* (one hour time steps) by the fraction of cells alive at time *t+1* over those cells alive at time *t.* For reproduction rates, we calculated the average number of divisions a cell underwent between time *t* and time *t+1* given that the cell was alive, this was done for each strain separately. The survival probabilities entered the sub-diagonal parameters of the strain specific Leslie matrix, and the age-specific division rates entered the top row of the strain specific Leslie matrix.

**Table 1:**
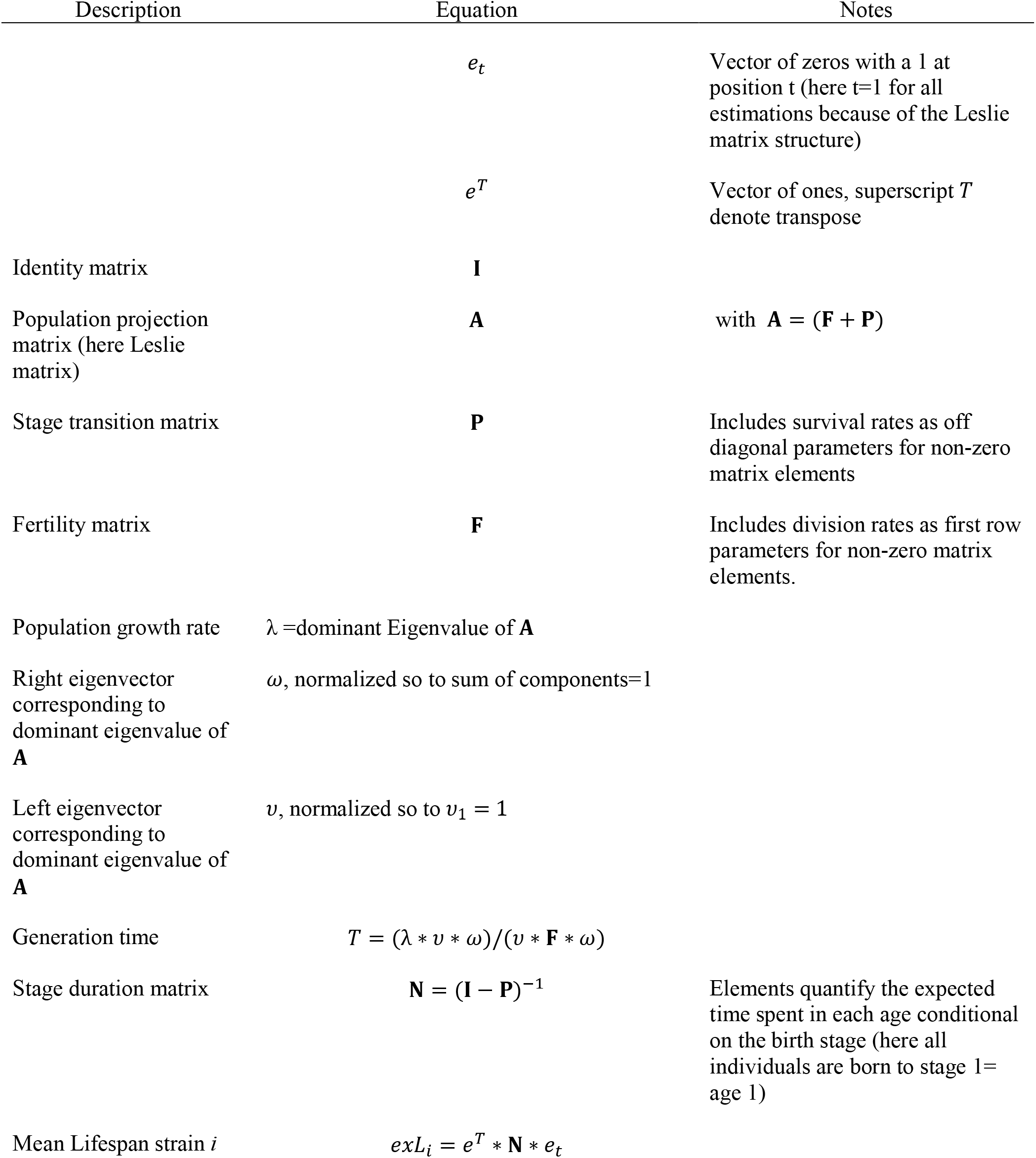
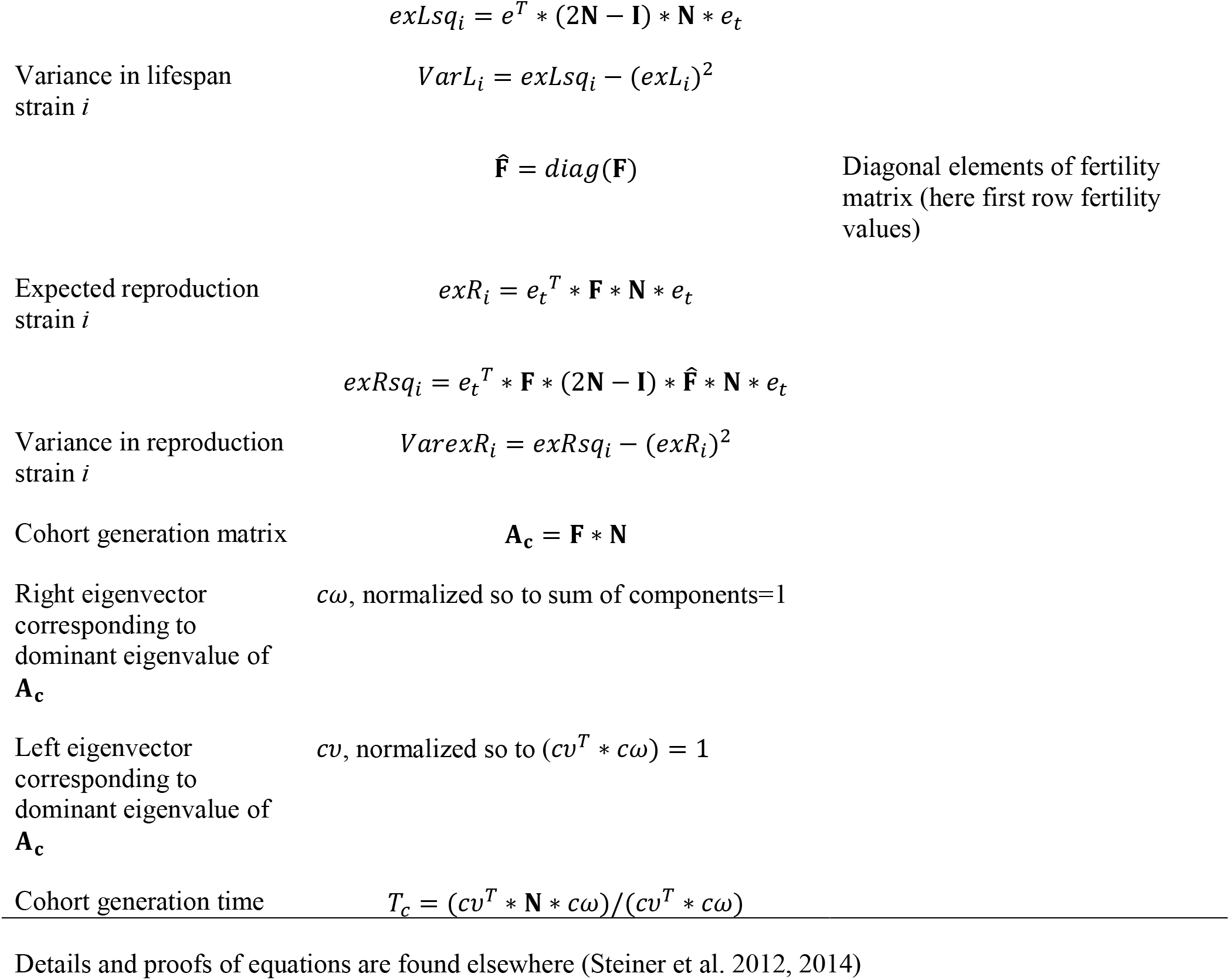
Notation and Equations

We choose a Leslie matrix model approach since these models conveniently and directly link individual level data, as collected by our experiments, to population level properties, including the population growth rate and the generation time. The direct calculation of vital rates, as commonly done for matrix models, rather than fitting function parameters as for instance done in logistic regressions, provided great variability and accuracy in estimating the demographic parameters. The close match between observed data and the matrix elements can be seen in Fig. A4. Such direct parameter calculation usually ignores the effect of sampling variability, and effects of sampling variability can be substantial for small populations (<100 individuals) with low survival (<0.5) (Fiske et al. 2008). In our study both survival rates and sample sizes (312 to 1017 cells per strain) were well above levels were substantial influences of sample variation is expected (Fiske et al. 2008). If such sampling variability would significantly influence our results, we would also expect to see low replicability among subsamples within strains, a pattern not found in our study (Fig. A3).

We assumed that all individual cells that were initially loaded into the microfluidic device are of age 0. We know that this assumption is partly violated. Based on stable-age theories of exponentially growing populations the age distribution is highly right skewed (Fig. A2), and less than 30% of the loaded cells are older than 1h, i.e. >70% of the individuals are of age<1h (Steiner et al. 2017).

Unfortunately, we have no means to determine which of the initially loaded cells are older than 1h. Convergence to a stable-age distribution, as assumed by matrix population models, should be fast in populations with vital rates as we computed for our bacteria populations. Further, such stable-age distribution should be closely achieved in the exponentially growing populations under constant and non-limiting growth conditions, as the ones the initially loaded mother cells are originating from. For the above reasons, potential transient dynamics are not expected to have large influences (SI Fig. A2) (Steiner et al. 2017).

We used the seven strain specific Leslie models, **A**, to compute for each strain the following demographic parameters: the population growth rate, λ, the cohort generation time, T_c_, the mean and among individual variance in lifespan, the mean and among individual variance in lifetime reproduction, the stable age distributions, and the age-specific reproductive values. Equations for estimating the demographic parameters are listed in Table 1, for proofs and further details please see Caswell (2001), Steiner and Tuljapurkar (2012), and Steiner et al. (2014). The computed demographic parameters are shown in Table 2, Fig. A1 and Fig. A2. We choose to estimate the mean and variance in fitness components — lifespan and reproduction — based on the Leslie matrix rather than on the original data to minimize the influence of different levels of right censoring. Fig. 3 shows the original observed data with the right censoring, and Fig. A4 shows the close match between the age at death distributions based on the original observed data and the age at death distribution predicted by the Leslie model.

**Table 2:** Key demographic parameters of the seven isogenic strains

| Strain | $\lambda$ | T | T<sub>C</sub> | Mean<br>Lifespan | SD<br>Lifesp. | CV<br>Lifesp. | Mean<br>LRS<br>(R<sub>0</sub>) | SD<br>LRS<br>(R<sub>0</sub>) | CV<br>LRS<br>(R<sub>0</sub>) | #<br>Individuals |
| --- | --- | --- | --- | --- | --- | --- | --- | --- | --- | --- |
| AB1157 | 2.02 | 2.50 | 31 | 23 | 28 | 1.2 | 42 | 54 | 1.3 | 504 |
| BW25113 | 2.25 | 2.34 | 77 | 54 | 24 | 0.4 | 128 | 57 | 0.4 | 482 |
| IAI1 | 1.74 | 3.29 | 69 | 44 | 31 | 0.7 | 74 | 56 | 0.8 | 312 |
| MG1655 Inlag | 2.24 | 2.36 | 43 | 34 | 26 | 0.8 | 72 | 57 | 0.8 | 519 |
| MG1655 LM | 2.18 | 2.30 | 44 | 34 | 26 | 0.8 | 67 | 54 | 0.8 | 505 |
| W3110 | 2.03 | 2.38 | 38 | 27 | 27 | 1.0 | 50 | 54 | 1.1 | 501 |
| MG1655 | 1.97 | 2.83 | 68 | 46 | 31 | 0.7 | 94 | 65 | 0.7 | 1017 |
|  |  |  |  |  |  |  |  |  |  | **3840** |

**Fig. 3:**
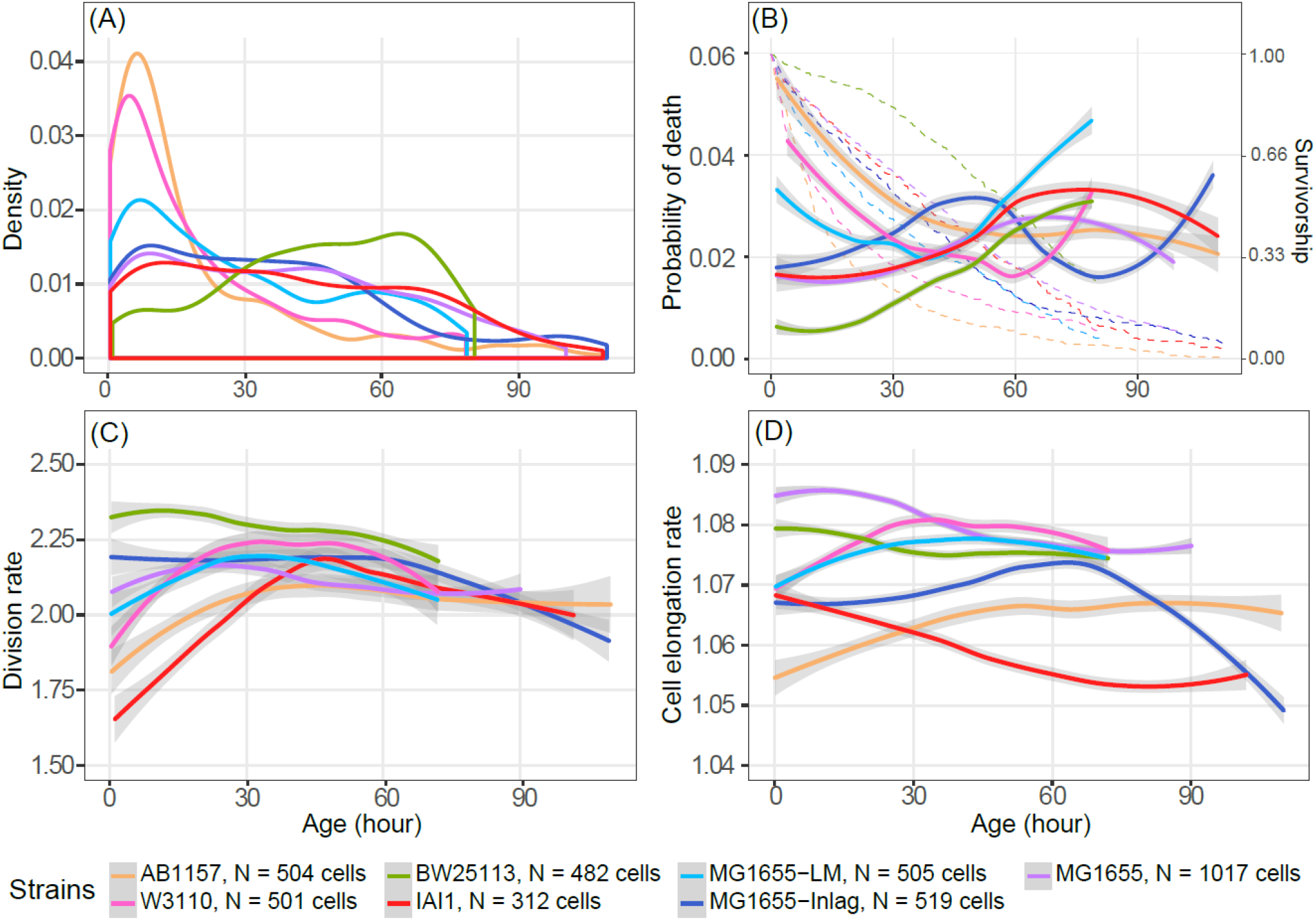
Lifespan distribution (A), probability of death (B), division rate (C), and cell elongation rate (D) of seven different bacteria strains plotted against age in hours. For (B) also survivorship curves are plotted as dashed lines. 95 % CI are shown in grey shading (B, C, D). For B and C hourly rates are shown, for D rates per 4min intervals are shown. Note, cells of the different strains are truncated (right censored) at different ages. All rates have been loess (program R) smoothed.

We decomposed the among strain variance (fixed genetic) and within strain (dynamic) variance in fitness components using the seven strain specific estimates of the variance in lifespan, *VarL*_*i*_ (equations: Table 1, values: Table 2; subscript *i* indicates the strain) and estimated the mean of these seven strain specific values. This mean variance 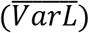 provided us with the mean within strain variance in lifespan. We followed the same procedure to compute the mean within strain variance in reproduction 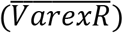. We related these mean (within strain) variances 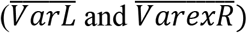 to the variance in the strain means 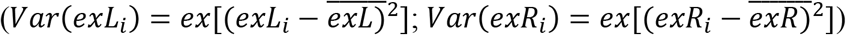 respectively. We equally weighted each strain estimate, i.e. we did not consider that some strains had more cells the estimates are based on compared to others. Our results are qualitatively robust to this assumption.

## Results

Our results are based on a total of 3840 individual cells (3461 were tracked over their whole lifespan and 379 were right censored) (Fig. 3A, B). The cells in our experiments originated from seven isogenic strains (312 to 1017 cells per strain; mean 549 ± 202 SD) (Table 2). Population growth rates, λ, varied between 1.74 and 2.25 per hour (mean 2.06 ± 0.17 SD), mean lifetime reproductive success (net reproductive rate, R_0_) varied, between 42 and 128 individuals (mean 75 ± 27), generation time, T, varied between 2.3 and 3.29 hours (mean 2.57 ± 0.34 SD), and cohort generation time, T_C_, varied between 31 and 77 hours (mean 52.9 ± 16.7 SD) (Table 2). The coefficient of variation in lifespan within strains (CV=within strain SD in lifespan/mean strain lifespan) varied between 0.4 and 1.2 (mean 0.8 ± 0.2), and was highly correlated to the CV of lifetime reproductive success within strains (0.4 to 1.3; across strain mean 0.8 ± 0.3). Mean within strain variance in lifespan was high at 766 h^2^ compared to variance in mean lifespan among strains at 105 h^2^. Similarly, mean within strain variance in lifetime reproductive success was high at 3230 ind^2^ compared to variance in mean lifetime reproductive success among strains at 708 ind^2^. Based on the variances within and among strains, ~88% of the variance in lifespan comes from within strains and 12% of variance in lifespan is caused by among strain variance. For lifetime reproductive success within strain variance dominates in generating ~82% of variance and 18% was observed among strains.

We illustrate the high variability in lifespan among individuals within strains in Fig. 3A. The corresponding age specific mortality patterns (Fig.3B) highlight the diversity in demographic patterns among strains. Such diversity is remarkable considering that all strains belong to the same species, *E. coli*, and have experienced identical constant environments throughout the experiments (highly controlled medium, nutrition, and temperature). Some strains showed negative chronological senescence with declining mortality with increasing age (AB1157), others showed more bathtub shapes with declining mortality early in life followed by classical senescence later in life (MG1655_LM, W3110), still others showed only classical senescence with increasing mortality with age (BW25113), or finally others first showed increased mortality early in life before exhibiting declining mortality later in life (MG1655, IAI1). Some of the late age mortality rates were estimated on small numbers of cells, and the very old age patterns (e.g. MG1655-Inlag showing steep rising mortality above 90h) should not be over-interpreted due to uncertainty from sampling variability. We also revealed substantial among strain diversity in age specific division rates (Fig. 3C) and cell elongation rates (cell growth rates) (Fig. 3D). Most strains reached somewhat similar division rates after age 45h and showed moderate decreases in division (reproductive senescence) at older ages. At younger ages division rates differed substantially among the different strains, by either being fairly constant or increasing with age. Cell elongation rates (cell growth grates) showed similar diverse patterns among strains as mortality. Cell elongation (cell growth) increased (e.g. AB1157), decreased (e.g. IAI1), or first increased and then decreased (e.g. MG1655-Inlag) with increasing age. The age-specific reproductive values and stable stage distributions for the seven different strains are shown in the Appendix (Supplementary material Appendix 1, Fig. A1, A2).

## Discussion

We showed that fixed heterogeneity in lifespan and reproduction, i.e. the genetic contribution, is moderate compared to the heterogeneity among individuals within strains, the neutral dynamic, and non-genetic and non-environmental, heterogeneity. Only our highly-controlled study system allows such an accurate and direct decomposition of heterogeneity in genetic and non-genetic contributions. Under less controlled setting, as in natural populations, we could not decompose the causes of mortality and reproduction without bias (Steiner and Tuljapurkar 2012, Bonnet and Postma 2016, Cam et al. 2016). Despite the dominating variability within strains, we detected significant and evolutionary important variability among strains. This selective difference is best illustrated by the differences in population growth rate, λ, which would lead to fast changes in genotype frequencies. The differences in λ directly inform us on each of the strains fitness, i.e. how fast the different strains would grow and compete against each other under the exponential growth conditions in our experiments. In our system, environmental conditions exclude any density dependence, reduce extrinsic environmental variability to a level that is negligible, and provide non-limiting conditions that promote exponential population growth as assumed under stable stage theories.

The within strain heterogeneity is partly illustrated by the coefficient of variation of the fitness components. The estimates we found here are comparable to less controlled systems and more complex organisms. In laboratory systems of other isogenic individuals under lab conditions the coefficient of variation (CV) ranges between 0.24 to 1.33 in lifespan [*Caenorhabditis elegans* 0.24-0.34 (Finch and Kirkwood 2000, Kirkwood et al. 2005), *Caenorhabditis briggsae* 0.31-0.51 (Schiemer 1982), *Saccharomyces cerevisiae* (0.37)(Kennedy 1994)]. Less genetically controlled lab populations do not differ much from these patterns in the CV of lifespan: laboratory reared mice (0.19-0.71)(Finch and Kirkwood 2000), *Drosophila melanogaster* 5.98-13.48 (Curtsinger et al. 1992). Even under less controlled conditions in the field, for instance, a plant species, *Plantago lanceolatum*, shows a CV 0.96 for lifespan, and 3.97 for reproduction (Steiner et al. unpublished) and such estimates seem not exceptional even in populations where we do not know the genetic or the environmental contributions (Tuljapurkar et al. 2009, Steiner et al. 2010). Even though in less controlled systems this CV includes contributions of fixed and dynamic heterogeneity, the comparatively similar estimates between highly controlled lab systems and natural systems might indicate that neutral variability could be substantial not only in controlled lab populations. If one sees it the other way around, our somewhat highly artificial and very simple model system shows surprisingly little difference in CV of fitness components compared to more natural systems.

The ambiguity of estimates in natural populations about fixed and dynamic heterogeneity generating observed variances has resulted in a heated debate about neutral and adaptive contributions to this heterogeneity (Bonnet and Postma 2016, Cam et al. 2016). As with other neutral theories in molecular biology (Leigh 2007), or community ecology (Hubbell 2001), the neutral theory of life histories (Steiner and Tuljapurkar 2012) has been attacked based on a common misunderstanding behind neutral theories, that is, the erroneous claim that all variability is neutral. We know that all neutral theories are wrong (Leigh 2007). Any natural population includes selective differences, but the neutral theory illustrates to what extend variability might be neutral (in its theoretical extreme). To date, we do not have the means to unambiguously differentiate the causes driving the observed heterogeneity in natural populations. In our study, we show how substantial neutral heterogeneity can be in a simple bacterial system. This large heterogeneity might be surprising since the strains are highly adapted to the lab conditions. Selection has not managed to get rid of this variability, and the question arises how such neutral variability is maintained and if it is adaptive.

Labeling heterogeneity among individuals as neutral is often perceived with skepticism, since apparent random processes might have a deterministic hidden biological cause. From a deterministic point of view, each individual would be born with an intrinsic clock that determines its life course. Such a clock would have no genetic or epigenetic component, because mother and daughters life course are not correlated (Steiner et al. 2017). We believe the bacteria system can partly inform on distinguishing among such a deterministic viewpoint and an understanding that explains these hidden underlying processes to be generated by random events showing, e.g. showing stochastic characteristics. Numerous molecular and biochemical processes that are assumed to shape life courses of individuals have been identified for *E. coli*, many of them related to direct or indirect oxidative processes, but as in any other system the molecular and biochemical process of aging for *E. coli* is not fully understood (Kirkwood et al. 2005, Raj and van Oudenaarden 2008, Lindner and Demarez 2009, Gómez 2010). Many of these mechanisms show in themselve stochastic properties, including e.g. stochastic gene expression, protein folding and misfolding, and their potential cascading effects up to the organism level (Elowitz et al. 2002, Lindner and Demarez 2009, Balázsi et al. 2011, Ackermann 2015). Despite these detailed insights on biochemical and molecular mechanisms that regulate intrinsic cellular processes, linking them to the individual life course remains challenging (Lindner et al. 2008, López-Otín et al. 2013). Such difficulties are expected if stochastic properties within the cells characterize these processes.

Our approach using Leslie models, only takes the age of the cell into account and averages individuals within the strains across traits, be they morphological (e.g. cell size), (stochastic) gene expression, or asymmetry in protein aggregates. Such averaging across trait variability should reduce the calculated variances in lifespan and reproduction among individuals belonging to the same strain, since stage dispersion in reproduction (among ages) caused by trait variability is reduced (Steiner et al. 2014). We could have included traits such as cell size, by extending our models to age-stage structured matrix models, formulated either as classical Lefkovitch matrixes or integral projection versions of matrix models (Caswell 2001, Ellner and Rees 2006). However, increasing the parameter space trades off against accuracy of parameter estimates due to sampling variability. Also, cell elongation and division rates are correlated in *E. coli* and therefore the Leslie matrices include — in the age dispersion in reproduction — part of the variability in cell size (Steiner et al. 2017). We aimed at a simple demographic model (Leslie matrix) that provides realistic and accurate estimates (Fig. A4). Extending these simple model to more complex models should be done in future studies. Among strain variance in fitness components should not be substantially influenced by averaging across individual trait variability. Obviously, we might have missed to explore important traits that are predominantly affected by the genetic differences among the strains. However, under our experimental conditions such traits, even if they had been highly differentiated among strains, did not have a significant influence either on reproduction or survival and therefore did not increase variability among strain fitness (λ). Using Leslie models directly linked the individual level data to population level properties without additional fitting of model parameters (Fig. A4). Fitting accurate functions to the somewhat complex demographic age patterns (Fig. 3) would have been challenging with other models. The Leslie matrix approach we choose provides accurate description at the population level, even though at older ages parameter calculation suffer from sampling variability. Such uncertainty at old ages should not strongly influence the overall variance decomposition, since the few individuals that live to old ages do not weigh heavily on the overall variance estimation.

We choose experimental conditions that reduced variance in certain traits, e.g. cell size. For instance we choose minimum medium M9 that reduces variance in division size (and size after division) compared to complex medium (Gangan and Athale 2017). Minimum medium M9 also decreases the rate of filamentation — a stress response where the cell continues to elongate without dividing. Under M9 conditions, filamentous cells rather died than recovering from filamenting by dividing into normal sized cells. Such recovery is frequently observed under complex media (Wang et al. 2010). Differences in experimental setup (e.g. starting with exponential or stationary growth cells, type of medium, strains explored, culturing devices used) make it challenging to direct compare to other single cell or batch culture *E. coli* studies, and even estimates within batch culture studies on growth rates are highly variable (Helmstetter 1968, Dennis and Bremer 2008). Compared to batch cultures grown on the same M9 media, our estimated exponential growth rates, λ, are high, though such increase in growth rates are expected and known for comparisons between single cell estimates and batch culture estimates that are in any case difficult to directly compare (Reshes et al. 2008) (Fig. A4).

In interpreting our results, we must be aware that all strains are subjected to some level of right censoring (mean: 1.4% to 26.3%; SD 9.3% ± 8.8). The number of individuals suffering from this censoring differs among the different strains and might therefore bias our results differently. We aimed at reducing the effect of the right censoring by estimating demographic parameters from Leslie matrices with open age brackets for the last age class (Fig. A4).

Another criticism on our data is that experiments are not entirely independently replicated. Each mother cell sits in its own little side channel, but the cells of each strain are still confounded in being loaded in the same microfluidic chip and have been provisioned by the same highly controlled laminar flow. The amount of nutrients delivered to the cells is magnitudes larger compared to the amount all cells could consume; hence there should be no limitation of resources or any difference in access to resources among cells. Preliminary experiments (Jouvet & Steiner unpublished) also indicate that diffusion properties among the individual side channels are very similar, ascertaining that extrinsic environmental variation the individual cell experience in the microfluidic device are negligible. Despite the confounding effects, we are convinced that our data is representative since patterns among independent flow channels are highly replicable as we illustrate in the SM for one of our strains (Supplementary material Appendix 1, Fig. A3). Further if individual side channels would differ in their environments we would expect a correlation between mother and daughter cells in their lifespan, but such correlation has not been found in other studies (Steiner et al. 2017).

Our results also illustrate how genetic variability, even within a species, can shape very diverse senescence patterns, both in survival and reproduction. Phylogenetically more closely related strains (Fig. 2) do not necessarily show more similar demographic patterns compared to less closely related strains (e.g. AB1157, MG1655, W3110). This raises interesting questions for comparative demography where a single population of a species is frequently assumed to be representative for each species (Jones et al. 2014). Even under our highly controlled environmental condition we see great diversity in demographic patterns and it would be interesting to compare multiple natural populations of the same species to investigate how persistent demographic patterns within species are in nature.

Given the highly controlled environment and the high genetic control our system also open doors to investigate basic evolutionary theories of life history (Hamilton 1966, Stearns 1992). Such theories base much of their arguments on a fundamental tradeoff between reproduction and survival or early versus late life trade-offs. One of the challenges of assessing such trade-offs include that individuals, populations, or genotypes receive different amounts of resources (energy or nutrients) and these differences might override the underlying trade-offs (van Noordwijk and de Jong 1986). Our highly controlled environment and the clear distinction of genotypes therefore provide a nice opportunity to reveal such tradeoffs that are hard to reveal in natural populations (Metcalf 2016). Based on the theories, we predict that strains with high mortality should exhibit high reproduction (high division and cell growth rates). Such simple expectations are not met, strains with relatively low mortality (e.g. BW25113) also showed high cell elongation and division rates, while other strains showed somewhat opposite patterns (e.g. AB1157). Similarly we did not detect clear age-specific trade-offs between early and late survival or early and late reproduction and their interaction as predicted by evolutionary theories of aging (Medawar 1952, Williams 1957, Hamilton 1966). Strain IAI1 for instance showed senescence in survival and in cell elongation rates, but increased in reproduction (division rate) with age, before plateauing off at old ages. Other strains (e.g. MG1655-Inlag) showed increased and decreased mortality with age and similar patterns in cell elongation, but did not show much change in division rate over much of life. Only late in life did MG1655-Inlag, reveal some reproductive senescence. We can interpret the lack of such expected relationships among survival and reproduction as a lack of genetic linkage between traits and ages, and that the underlying life-history tradeoffs are not as strong as assumed. One might argue that this system is too artificial to express such trade-offs.

Still, we see familiar demographic patterns even in this simple system, and our best evidence for such trade-offs is coming from such artificial lab organisms, rather than from populations in their natural environments (Metcalf 2016). A fundamental challenge behind revealing these trade-offs is that they are expressed within individuals and not among individuals, but most of our attempts compare among individuals that belong to different groups, genotypes, populations, and species, as we do in our study.

Our findings unambiguously quantified fixed and dynamic heterogeneity for a simple bacterial system. We revealed that substantial variability is generated by cell-intrinsic likely stochastic processes and that the quantity and timing of these processes differ among the clonal strains, shaping diverse age-specific demographic patterns. To what extent similar levels of variability are generated by intrinsic likely stochastic processes in natural populations of simple organisms such as bacteria or more complex organisms should be explored. We discussed similarities in coefficient of variation across different level in complexity among organisms and across levels of control that suggest that our result is not exceptional. Promising attempts to overcome the unknown genetics of individuals in natural populations have been made by releasing hundreds or thousands of genetically known crossed individuals into the wild and then tracked throughout their lives (Roach 2012, Travis et al. 2014). Evidence of such experiments suggests that levels of within cross heterogeneity is substantial compared to among cross heterogeneity. How such heterogeneity is maintained, how it has evolved, and whether it is adaptive remains to be explored.

## Acknowledgements

We thank all members of the Max Planck Odense Center on the Biodemography of aging for discussions and comments. We were supported by the Max Planck Society (LJ, UKS) and SFB 973 (Deutsche Forschungsgemeinschaft), project C5 (ARR).

